# Cerebellar climbing fibers signal flexible, rapidly adapting reward predictions

**DOI:** 10.1101/2024.10.09.617467

**Authors:** Carlo Vignali, Michael Mutersbaugh, Court Hull

## Abstract

Classical models of cerebellar computation posit that climbing fibers (CFs) operate according to supervised learning rules, correcting movements by signaling the occurrence of motor errors. However, recent findings suggest that in some behaviors, CF activity can exhibit features that resemble the instructional signals necessary for reinforcement learning, namely reward prediction errors (rPEs). Despite these initial observations, many key properties of reward-related CF responses remain unclear, thus limiting our understanding of how they operate to guide cerebellar learning. Here, we have measured the postsynaptic responses of CFs onto cerebellar Purkinje cells using two-photon calcium imaging to test how they respond to learned stimuli that either do or do not predict reward. We find that CFs can develop generalized responses to similar cues of the same modality, regardless of whether they are reward predictive. However, this generalization depends on temporal context, and does not extend across sensory modalities. Further, learned CF responses are flexible, and can be rapidly updated according to new reward contingencies. Together these results suggest that CFs can generate learned, reward-predictive responses that flexibly adapt to the current environment in a context-sensitive manner.

## Introduction

Cerebellar climbing fibers (CFs) are thought to play a key role in instructing associative learning (Albus, 1971; Ito, 1972; Marr, 1969). To mediate such learning, CFs convey error signals to the cerebellar cortex based on sensorimotor feedback (Medina et al., 2000). Classical models posit that these error signals follow the principles of supervised learning, a framework that is supported by measured CF activity in many cerebellar-dependent behaviors (Raymond and Medina, 2018). However, recent work has also revealed features of CF signaling that diverge from supervised learning principles (Hull, 2020). Across multiple behaviors guided by the anticipation of reward, evidence has emerged that CFs can signal predictions, and evaluations of those predictions, about reward delivery (Heffley and Hull, 2019; Heffley et al., 2018; Kostadinov et al., 2019; Larry et al., 2019; Sendhilnathan et al., 2021). These reward-related CF responses match some of the predictions established by the principles of reinforcement learning, and suggest the possibility that CFs signal a form of reward prediction error (rPE) to modify behavior in certain tasks (Hull, 2020; Kostadinov and Hausser, 2022). However, many features of reward signaling remain to be tested for CFs, and some aspects of CF activity, such as the apparent absence of negative prediction errors, do not match canonical properties of rPE signals^7^. Therefore, the essential features of cerebellar CF reward signals, and how they operate to promote learning and behavior, have remained unclear.

To appropriately guide learning, rPE signals must distinguish sensory cues according to the information they provide about reward (Sutton and Barto, 1998). While several types of rPE signals have been proposed (Bromberg-Martin et al., 2010), leading models of reinforcement learning typically require that these signals should scale with cue-associated reward expectation, and represent expected reward value with both increases and decreases in neuronal activity to indicate positive and negative value predictions (Rescorla and Wagner, 1972; Sutton and Barto, 1990). Other forms of rPE signals, however, have been suggested to follow somewhat different rules. For example, some forms of rPE can signal value with unidirectional changes in neural activity, behaving as so-called "unsigned" prediction errors (Pearce and Hall, 1980). Regardless of the type of signal, rPEs should at minimum distinguish cues that predict reward from other sensory input. This basic property is a key feature that delineates rPE signals from other sensory driven responses such as those associated with salience or novelty (Tapper and Molas, 2020) (though such signals may also play a role in reinforcement learning (Kakade and Dayan, 2002; Laurent, 2008)). In addition, rPE signals must also be flexible, and update their responses to reflect changing environmental reward contingencies (Tobler et al., 2005).

To begin understanding how the instructional signals carried by cerebellar CFs operate in reward-based tasks, we have tested these two key properties by evaluating their response to sensory stimuli of different modalities that either do or do not predict reward. Specifically, we have used a Pavlovian conditioning task to measure learned, reward-predictive CF responses, and tested how they depend on the similarity of the cues and the temporal structure of the task. Surprisingly, for similar cues of the same sensory modality, we find that CFs can produce equivalent learned responses if the cues are presented in close temporal proximity, even though mice expertly discriminate the cues behaviorally. This generalization, however, is much weaker if the cues are separated in time. Moreover, generalization does not extend across modalities. These results suggest that learned CF responses are sensitive to context, and do not simply reflect sensory input that predicts reward or training history. To further probe the flexibility of these responses, we reversed the stimulus-reward contingencies in trained animals. These experiments revealed that CFs can rapidly modify their responses, updating CF signals to reflect current reward-predictive sensory input. Together, these data reveal that cerebellar CFs can flexibly encode reward-predictive sensory input based on current conditions in a manner that is consistent with many features of reward-based learning systems.

## Results

### Mice can expertly discriminate visual cues that differentially predict reward

To examine how climbing fibers (CFs) represent two conditioned stimuli with different reward associations, we used a modified classical conditioning task in combination with two-photon (2P) imaging of cerebellar Purkinje cell dendrites (Heffley and Hull, 2019). In this task, mice learned to discriminate between two visual gratings presented randomly in time that differ according to their orientation and drift direction, with one stimulus predicting reward delivery (Vis+; horizontal, upward drifting) and one that does not (Vis-; vertical, rightward drifting) (**Fig. 1A,B**). After training (learning = 28.5 ± 4.8 sessions, **Fig 1C**), mice learned to accurately discriminate between these cues, licking reliably only in response to Vis+. Learning included both a significant increase in hit rates (pre-learning, 0.37 ± 0.10; post-learning, 0.95 ± 0.03; p=0.001, paired t-test, n=6 mice, **Fig. 1D**) and correct reject rates to Vis-(pre-learning, 0.65 ± 0.08; post-learning, 0.94 ± 0.03; p=0.03, paired t-test, **Fig. 1D**). We also observed a trend toward decreased reaction time on Vis+ trials across learning (pre-learning, 947.6 ± 59.7 msec; post-learning, 896.0 ± 55.8 msec; p=0.61, paired t-test, **Fig. 1E**), though this measure is influenced by the strategy of stochastic licking adopted by naive mice to continuously sample for reward. Overall, these data indicate that mice successfully learned to discriminate between two similar visual cues, exhibiting little or no behavioral generalization to Vis-(**Fig. 1F,G**).

**Figure 1.**
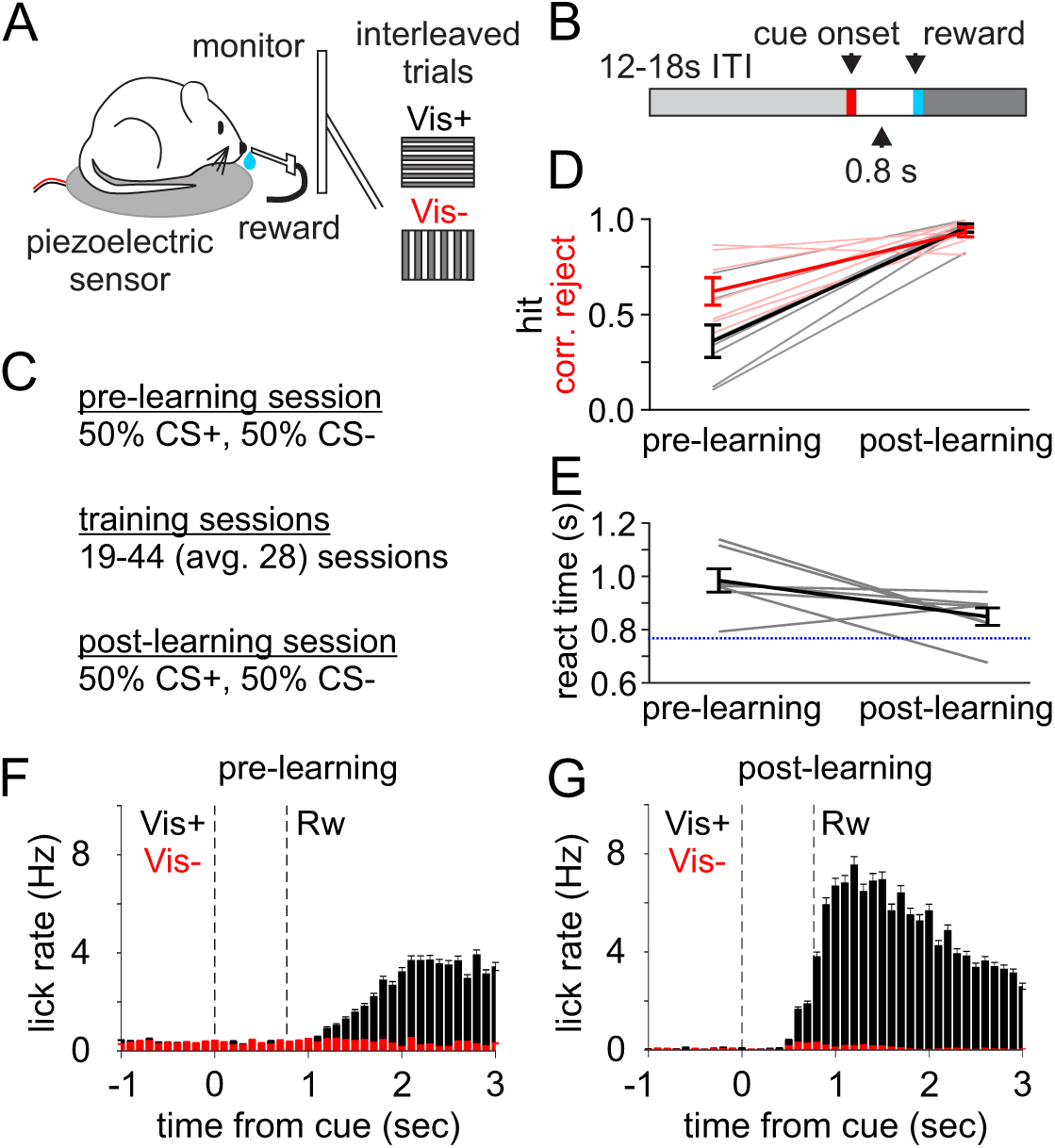
Pavlovian task to associate reward predictive and non-reward predictive visual cues. **A)** Schematic of experimental design. Mice viewed a grey screen and were transiently presented with one of two moving gratings for 767 ms. Reward delivery was synchronized with the termination of Vis+ cue presentation. **B)** Schematic of trial design. ITI-inter-trial interval. **C)** progression of training. **D)** Summary of hit (black) and correct reject (red) rates before and after learning. **E)** Summary of reaction time to reward delivery on Vis+ trials (dashed line indicates reward delivery). **F,G)** Summary histograms of mean lick rates across experiments before (**F**) and after (**G**) learning. Histograms are aligned to cue onset at time 0 and reward delivery (Rw) 767ms after cue onset. All data are presented as mean +/- SEM.

### CF activity is associated with licking in naive mice

To test how CFs encode Vis+ and Vis-, we performed 2P imaging in lobule simplex, an area that exhibits reward-related activity across multiple behaviors (Heffley and Hull, 2019; Heffley et al., 2018; Kostadinov et al., 2019; Wagner et al., 2017). By expressing GCaMP7f in Purkinje cells using a Cre-dependent strategy, we measured the postsynaptic input of CFs to PC dendrites, so called "complex spikes" (Cspks, see **Methods**)(Heffley and Hull, 2019; Heffley et al., 2018). We first compared the rate of CF inputs in naive mice following cue and reward delivery. While we did not observe a well-timed CF response to visual input or reward delivery, we did observe a slow increase in CF activity during the trial that continued through reward delivery on both Vis+ (mean = 1.04±0.03 Hz; p=9.5e-22, paired t-test) and Vis-trials (mean = 1.15±0.04 Hz; p=1.5e-25, paired t-test; **Fig 2A**). In agreement with prior work (Heffley and Hull, 2019), this increase in CF input is associated with the first lick during the trial, and can therefore be observed most readily by aligning Cspks to the first lick (Vis+ = 2.01±0.07 Hz; p=6.5e-55, paired t-test; n=262/358 trials, 433 cells; 6 mice; Vis-= 2.23±0.09 Hz; p=5.1e-53, paired t-test; n=150/308 trials, 433 cells; 6 mice; **Fig. 2A, inset**). Because these responses occur on both Vis+ and Vis-trials, they may be related to either anticipation or motor output, but not the detection or consumption of reward.

**Figure 2.**
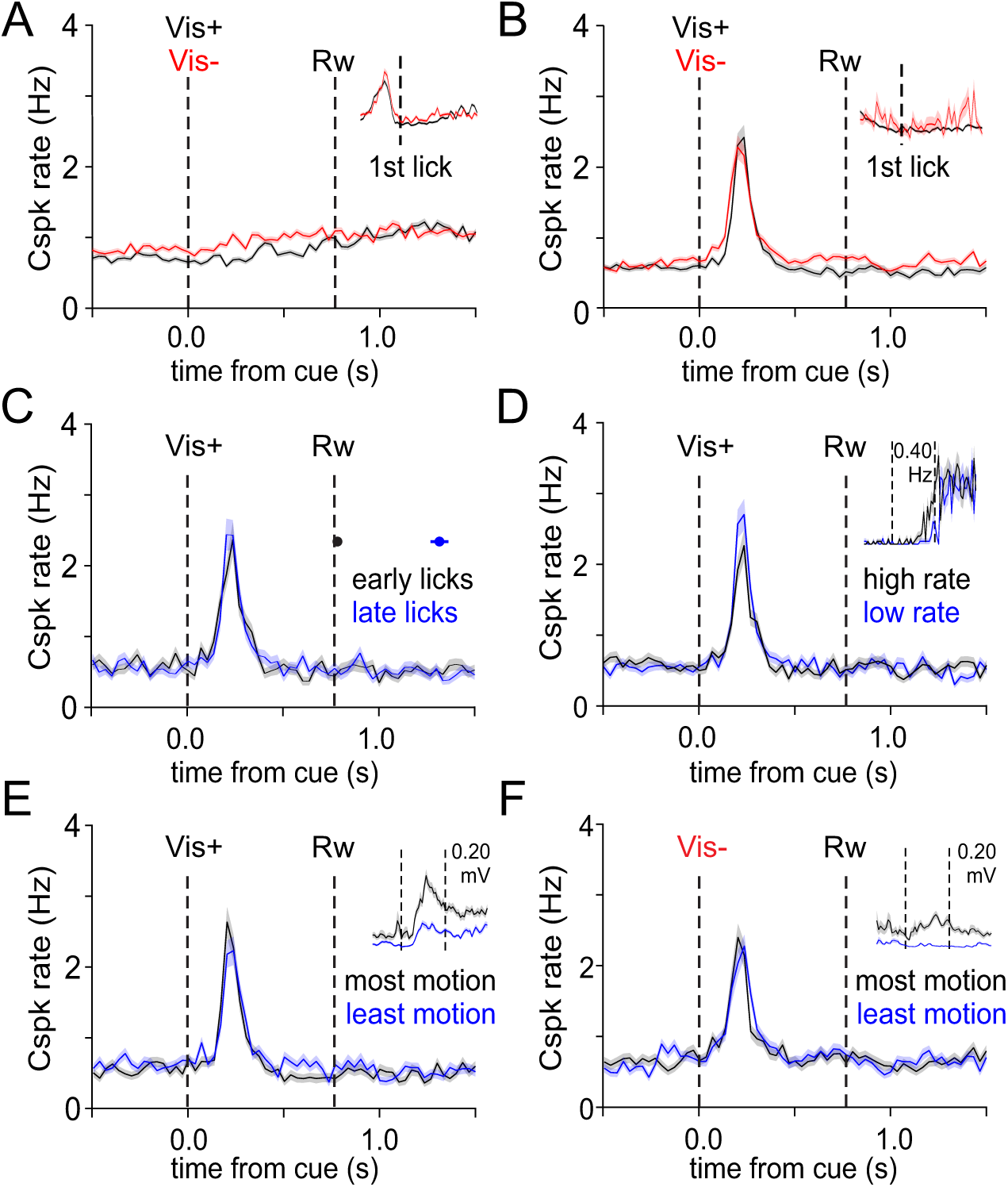
CFs respond to both Vis+ and Vis-. **A)** Peristimulus time histogram (PSTH) of complex spike (Cspk) rate aligned to cue onset on Vis+ (black) and Vis-(red) trials in naive animals. Dashed lines indicate cue onset and cue offset, coincident with reward delivery on Vis+ trials. Inset, Cspk rate aligned to the first lick after cue presentation in naive mice. Lick- and cue-aligned plots share the same y-axis scale. Lick aligned plots show 500 ms before through 1000 ms after lick onset; dashed line represents lick onset. **B)** Same as A) but for post-learning. **C)** PSTH of post-learning Cspk rates on Vis+ trials with the earliest (black) and latest (blue) licks. Dots represent mean onsets for earliest and latest 20% of lick bursts. **D)** Same as C) for trials with the highest (black) and lowest (blue) rates of pre-reward licking. Inset, PSTH of mean lick rates. **E,F)** Same as C) for trials with the most (black) and least (blue) animal motion on Vis+ (**E**) and Vis-(**F**) trials. Insets, PSTHs of mean piezo voltage traces. All data are presented as mean +/- SEM.

### CFs generalize in response to visual stimuli with different reward associations

Consistent with previous work (Heffley and Hull, 2019), we find that CFs develop robust responses to the reward predictive visual stimulus after learning, (Vis+: mean = 2.08 ± 0.12 Hz; p=6.8e-33, paired t-test; n = 316 neurons, 6 mice; **Fig. 2B**). Surprisingly, however, CFs also developed a learned response to the non-reward predictive Vis-stimulus, even though mice were experts at discriminating between the cues (mean = 2.06 ± 0.13 Hz; p=1.9e-24, paired t-test; 316 neurons; **Fig. 2B**). In fact, we observed no difference in peak CF response amplitude to the two stimuli (p=0.49, Wilcoxon Ranked Sum Test). These data reveal that under these stimulus conditions CFs generalize responses despite behavioral discrimination. At the level of individual CF inputs, we did not observe unique responses to Vis+ or Vis-. Approximately 1/3 of CFs responded to both Vis+ and Vis-(112/316 cells). Of the remaining cells, a lower fraction preferentially responded to either cue, with more cells responding to Vis+ than Vis-(Vis+ = 79 cells and Vis-= 32 cells; **Supp. Fig. 1A,B**). Thus, the same CFs can exhibit learned responses to both reward predicative and non-reward predictive cues.

After training, as with previous experiments (Heffley and Hull, 2019) we found that CF lick-aligned responses were abolished (**Fig. 2B, inset**), suggesting that CF activity does not represent motor output or lick-associated reward anticipation in trained animals. To further test this hypothesis, we measured how CF responses varied with lick timing and lick rate. We observed no difference in the latency or amplitude of CF responses associated with lick timing (Earliest 20% licks: CF latency= 134.97 ± 6.05 ms; CF amplitude = 2.39 ± 0.24 Hz; Latest 20% licks: CF latency 124.05 ± 5.91ms; CF amplitude = 2.45 ± 0.21 Hz, latency p=0.24, amplitude p=0.78, paired t-tests; **Fig. 2C**). Similarly, CF responses did not depend on lick rate (Lowest 20% lick rate: CF response = 2.73 ± 0.22 Hz; Highest 20% licks: CF response = 2.45 ± 0.22 Hz; p=0.12, paired t-test; **Fig. 2D**). Importantly, animals did not consistently lick during Vis-trials, thus ruling out licking as a primary driver of neural activity in these trials (**Fig. 1G**).

We also considered whether CF responses may reflect other body movements, as mice can exhibit anticipatory movements following the sensory cues after learning. We therefore segregating trials according to animal movement (Heffley and Hull, 2019), but found no significant differences in CF responses during either Vis+ or Vis-trials related to movement (Vis+ least movement = 2.34 ± 0.22 Hz; Vis+ most movement = 2.29 ± 0.20 Hz; p=0.82, paired t-test; **Fig. 2E**; Vis-least movement = 2.28 ± 0.15 Hz; Vis-most movement = 2.43 ± 0.20 Hz; p=0.15, paired t-test, **Fig. 2F**). Together these data suggest that the post-learning CF responses are associated with the visual cues and not motor output related to licking or body movement.

### Learned CF responses are sensitive to reward context

What could be responsible for the generalized CF responses to both Vis+ and Vis-? In the ventral tegmental area (VTA), dopamine neurons can also develop generalized responses to unrewarded stimuli under specific circumstances (Kobayashi and Schultz, 2014). These effects have been attributed to reward context, and shown to depend on the similarity of the cues and their temporal association with reward delivery. Accordingly, generalization is strongest when conditioned stimuli are similar and share a common sensory modality, and when they are presented on randomly interleaved trials that places both stimuli in relatively close temporal association with reward (Kobayashi and Schultz, 2014; Nakahara et al., 2004).

To test the role of context in learned CF responses, we first segregated Vis+ and Vis-in time by presenting them in separate trial blocks (**Fig. 3A**). In the same animals that had previously experienced interleaved Vis+ and Vis-trials, we found that CF responses were larger on Vis+ trials when the stimuli were presented in trial blocks (Vis+ = 1.79 ± 0.03 Hz; n=380 neurons; Vis-= 1.24 ± 0.02 Hz; n = 369 neurons, p=0.001, Wilcoxon ranked sum test; 4 mice; **Fig. 3B**). The larger response to Vis+ did not depend on the order of trial blocks, as CF responses were larger when the Vis+ block was presented first (Vis+ = 1.51 ± 0.12Hz; Vis-= 1.18 ± 0.08Hz; p=0.03, Wilcoxon ranked sum test; Fig 3B, top) or second (Vis+ = 2.10 ± 0.15Hz; Vis-= 1.36 ± 0.12Hz; p=0.03, Wilcoxon ranked sum test, **Supp. Fig 2A,B**). These data suggest that, like VTA dopamine neurons, learned CF responses are sensitive to trial structure and the temporal relationship between cues and reward.

**Figure 3.**
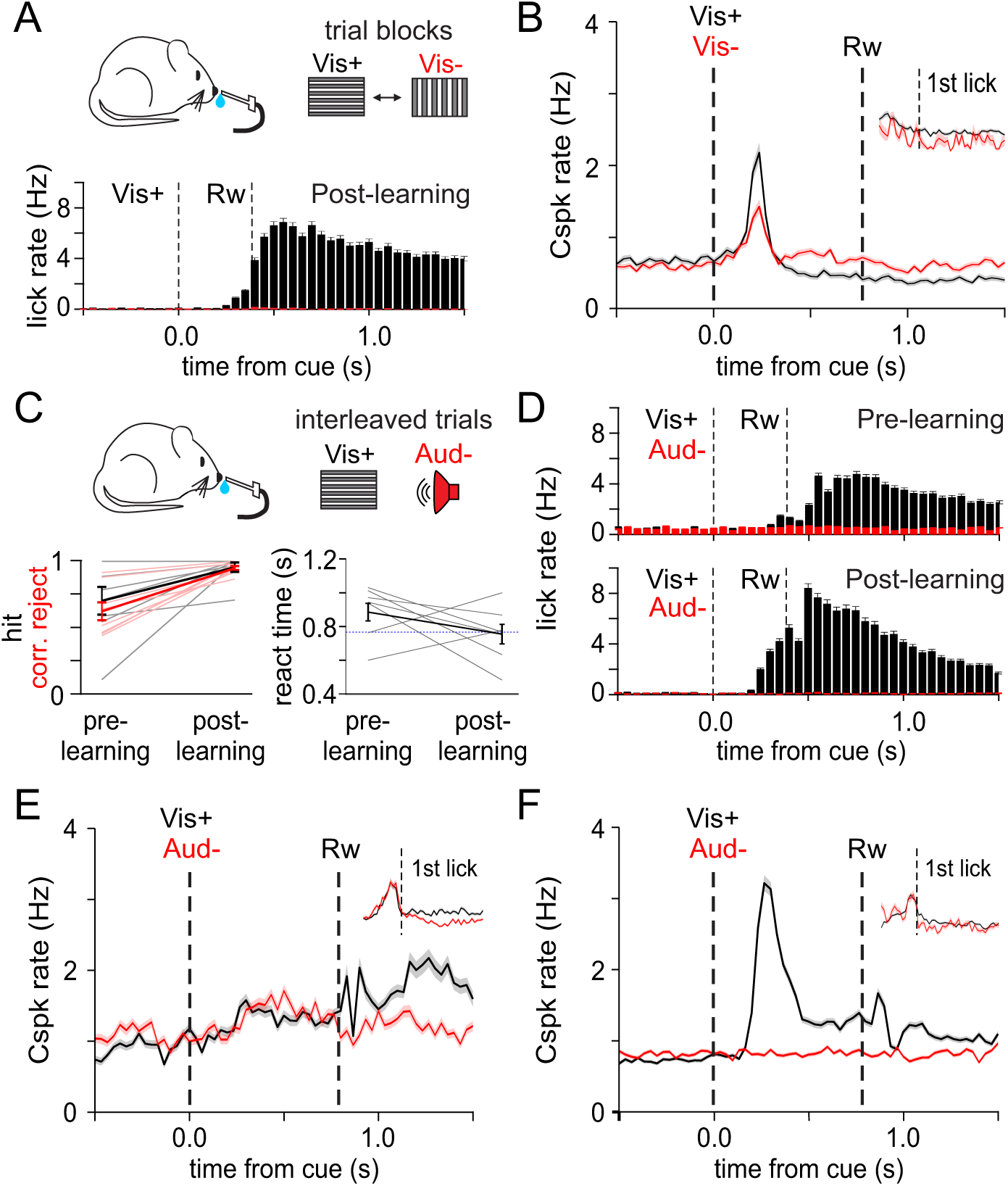
CF responses depend on trial structure and stimulus similarity. **A)** Top, Schematic of task design, with Vis+ and Vis-presented in blocks of 60 trials. Bottom, mean lick rates histograms for blocked sessions. **B)** PSTH of Cspk rates aligned to cue onset during block presentation of Vis+ (black) and Vis-(red). Dashed lines indicate cue onset and cue offset, respectively, coincident with reward delivery on Vis+ trials. Inset, Cspk rate aligned to first lick after cue presentation. Lick- and cue-aligned plots share the same y-axis scale. **C)** Top, schematic of task design for random presentation of Vis+ (black) and Aud-(red). Bottom left, summary of hit (black) and correct reject (red) rates. Bottom right, reaction time to reward delivery on Vis+ trials. Dashed line indicates reward delivery. **D)** Mean lick rate histograms for pre- and post-learning sessions. **E,F)** Same as B) but for pre-(**E**) and post-learning (**F**) sessions with Vis+/Aud-trials aligned to cue presentation. Inset shows alignment to first lick following cue presentation. All data are presented as mean +/- SEM.

To further test the requirements of learned CF responses, we made the reward-predictive cues more distinct by placing them in separate modalities. Specifically, we trained mice using a visual cue (Vis+; horizontal, upward drifting) that predicted reward, and an auditory cue (Aud-; tone kHz/dB/ms) that did not (**Fig. 3C**). As in the visual-only task, mice learned to expertly discriminate these two stimuli, licking reliably only in response to Vis+ after learning (**Fig 3C,D**). Accordingly, learning was associated with a significant increase in both hit rate (Vis+: 0.70 ± 0.10 → 0.96 ± 0.04; p = 0.04, paired t-test, **Fig. 3C**) and correct reject rate (Aud-: 0.62 ± 0.07 → 0.95 ± 0.02; p = 0.002, paired t-test, **Fig. 3C**), as well as a trend towards decreased reaction time across learning during Vis+ trials (pre-learning 1,028.9 ± 62.1 msec; post-learning, 813.1 ± 88.1 msec; p=0.14, paired t-test, **Fig. 3C,D**).

In these experiments, following learning CFs again generated a robust learned response to Vis+ (mean = 3.00 ± 0.08 Hz; p=1.2e-112, paired t-test; n = 707 neurons, 8 mice, **Fig. 3 E,F**). However, CFs did not develop a learned response to Aud-(mean = 0.86 ± 0.02 Hz; p=0.61, paired t-test; 707 neurons, 8 mice; **Fig. 3F**). Interestingly, CFs also maintained a smaller but measurable response to the first lick following Vis+ after learning (mean = 2.24 ± 0.08 Hz; p=5.2e-69, paired t-test; 185/707 cells) and Aud-(mean = 2.18 ± 0.10 Hz; p=2.2e-42, paired t-test; 87/707 cells; **Fig 3F, inset**). This is likely explained by the much more rapid time to proficiency in this task, which was only 12 ± 1.2 sessions, compared with 28.5 ± 4.8 sessions in Vis+/Vis-experiments. Overall, these data reveal that, like midbrain dopamine neurons, generalization of CF responses also depends on the similarity of sensory cues.

### Mice flexibly relearn reward contingencies

Our data suggest that CF responses can be generalized based on reward context and stimulus similarity. However, the absence of learned responses to auditory stimuli may not be due to a lack of generalization, but instead an inability for CFs to represent this modality in lobule simplex. Yet another possibility is that learned CF responses depend on training history. For example, during Vis+/Vis-learning, mice begin licking to both stimuli early in training before learning to disregard Vis-. This early behavioral generalization did not occur for Vis+/Aud-training. Therefore, CF responses could be determined by behavioral generalization early in training, and thus reflect only the first stimulus or stimuli perceived to be associated with reward. To test the flexibility of learned CF responses to different sensory modalities and training contingencies, we reversed the reward contingencies in the same mice after they had successfully learned to discriminate Vis+ and Aud-(**Fig. 4A**). After reversal, mice quickly learned the new contingency (learning = 10.3 ± 0.6 sessions), licking reliably only in response to the Aud+ (Correct rejects: reversal day = 0.22 ± 0.10; post-learning, 0.82 ± 0.04; p=0.006, Hit rates: reversal day = 0.70 ± 0.15; post-learning = 0.93 ± 0.04; p=0.22, paired-test, n=5 mice, **Fig. 4B-D**).

**Figure 4.**
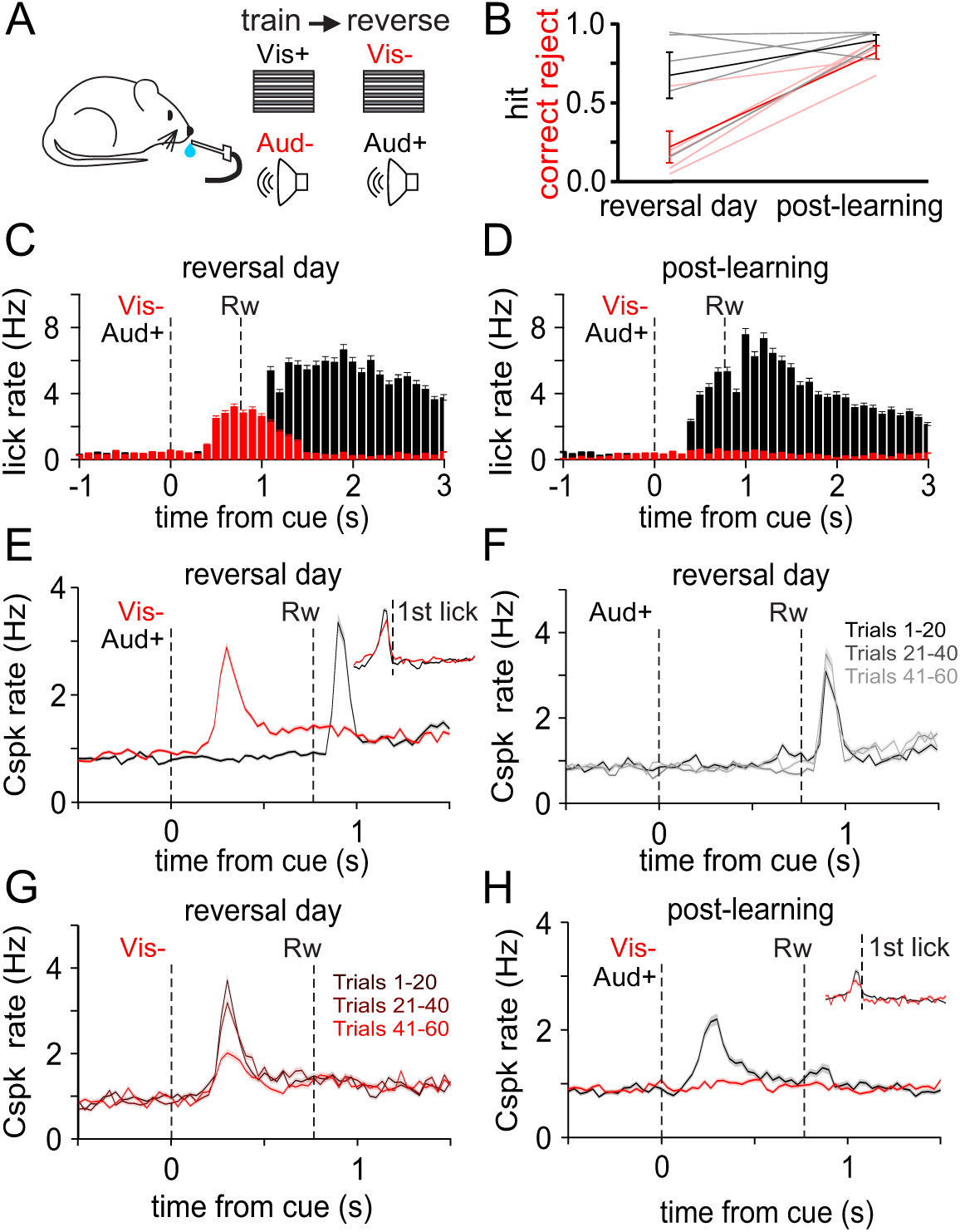
CF responses can be flexibly modified by new learning. **A)** Schematic of task design for reversal learning sessions. **B)** Summary of hit (black) and correct reject (red) rates before and after learning. **C,D)** Mean lick rate histograms for Aud+ (black) and Vis-(red) trials before (**C**) and after (**D**) reversal learning. **E)** PSTH of Cspk rates on reversal day aligned to cue onset and first lick following Aud+ (black) and Vis-(red) presentation. Inset, Lick- and cue-aligned plots share the same y-axis scale. **F,G)** PSTH of Cspk rates from E) segregated into three trial epochs for Aud+ (**F**) and Vis-(**G**) trials, respectively. **H)** PSTH of Cspk rates during post-learning sessions and lick-aligned PSTH (inset). All data are presented as mean +/- SEM.

### CF rapidly update predictive value of cues as reward association changes

On the first day of contingency reversal, we find that CFs maintained a robust response to Vis-(mean = 2.55 ± 0.07 Hz; p=3.8e-90, paired t-test; 498 cells, 5 mice, **Fig. 4E**). Moreover, we observed no response to Aud+ (mean = 0.84 ± 0.02 Hz; p=0.28, paired t-test; 498 cells, 5 mice; **Fig. 4E**). We also observed large CF responses to the first lick following both Aud+ (mean = 3.49 ± 0.10 Hz; p=3.3e-107, paired t-test; 498 cells, **Fig. 4E**) and Vis-(mean = 2.84 ± 0.09 Hz; p=2.6e-76, paired t-test; n=498 cells; **Fig. 4E**). This response was evident on Aud+ trials even without aligning to licks, as mice were already experienced with the task format and executed licks immediately following reward delivery. The re-emergence of large lick responses to both trial types may reflect defied reward expectations, as such responses were consistently abolished when reward was fully expected (Heffley and Hull, 2019) (**Figs. 2B,3B**).

To test whether these CF signals can flexibly represent changing task conditions, we first examined how they changed within the first session after contingency reversal. Here, we found that the response to ‘unexpected’ reward on Aud+ trials remained relatively stable, exhibiting a small increase across trials (first third: mean = 2.31 ± 0.13 Hz; middle third: mean = 2.73 ± 0.15 Hz; last third: mean = 2.87 ± 0.15 Hz; p=1.6e-13, repeated measures ANOVA; **Fig. 4F**). We also did not observe a CF response following Aud+ delivery on the first day of contingency reversal (first third: mean = 0.79 ± 0.05 Hz; middle third: mean = 0.77 ± 0.05 Hz; last third: mean = 0.79 ± 0.05 Hz; p=0.87, repeated measures ANOVA; **Fig. 4F**). However, we did observe a large decrease in the response to Vis-across trials (first third: mean = 3.12 ± 0.13 Hz; middle third: mean = 2.85 ± 0.12 Hz; last third: mean = 1.83 ± 0.09 Hz; p=3.4e-67, repeated measures ANOVA; **Fig. 4G**). Hence, these data revealed a rapid transition of CF responses away from the stimulus that was no longer reward predictive.

After learning, we observed a dramatic change in CF responses. Specifically, we found that CFs produced a large response to Aud+ (mean = 2.03 ± 0.08 Hz; p=7.3e-37, paired t-test; n =397 cells, 5 mice; **Fig. 4H**) that did not occur in response to Vis-(mean = 1.04 ± 0.03 Hz; p=2.1e-34, paired t-test vs Aud+ response; **Fig. 4H**). These data thus demonstrate a complete reversal of CF activity to reflect a newly learned reward contingency. Notably, we found that CFs still responded somewhat to the first lick on trials of both types, though the responses were significantly decreased as compared with the contingency reversal day (Aud+: p=8.43e-16; Vis-: p=2.30e-11; two-sample t-test, **Fig. 4H**). This may again reflect the much shorter training period (∼10 sessions) for this learning timepoint. Overall, these data demonstrate that learned CF responses are flexible, and can be readily modified to reflect new reward associations.

## Discussion

We used a modified Pavlovian conditioning task to test whether climbing fibers (CF) in lobule simplex differentiate between stimuli that do and do not predict reward. By measuring CF responses before and after learning, we found that CFs generate learned responses that depend on reward context, and preferentially represent reward-associated stimuli under conditions that segregate stimulus-reward associations. In addition, we find that learned CF responses are not fixed by training history, but instead can be flexibly modified to represent changing stimulus-reward contingencies.

In line with prior work, we find that CFs develop learned responses to reward-predictive conditioned stimuli (Heffley and Hull, 2019; Kostadinov et al., 2019). However, previous work had only assessed CF responses to a single stimulus that was reward-predictive^7^. Here, we find that CFs can develop learned responses to multiple sensory inputs, including those of different sensory modalities. Surprisingly, however, these learned responses are not exclusive to stimuli that predict reward in all contexts. Instead, we find that when conditioned stimuli are similar, sharing the same sensory modality and presented in close temporal proximity, CFs develop learned responses to both rewarded and unrewarded stimuli. In this way, CFs can develop generalized responses that are not exclusively yoked to the expectations of reward associated with specific stimuli or immediate task performance.

While these generalized CF responses are not typically predicted by leading models of reinforcement learning (Glimcher, 2011; Keiflin and Janak, 2015), they are consistent with the context-sensitive responses of ventral tegmental area dopaminergic (DA) neurons (Kobayashi and Schultz, 2014; Nakahara et al., 2004; Schultz, 2010). There, DA neurons develop responses to unrewarded stimuli in contexts where reward is predicted by another stimulus, even when an animal expertly distinguishes the rewarded and unrewarded stimuli at the behavioral level. Moreover, similar to the CF responses we have measured, reducing contextual similarity produces differential DA neuron responses to rewarded and unrewarded stimuli. While the utility of such generalization has not been resolved, it has been suggested to promote reinforcement learning by encouraging appropriate higher-order learning to potentially rewarding stimuli (second-order conditioning)(Kobayashi and Schultz, 2014). In the case of cerebellar learning, these data indicate that learned CF responses cannot be assumed to simply map onto learned stimuli that predict reward, or that they will provide a clear readout of immediately upcoming behavior.

Our data also support a key difference from the responses of many midbrain DA neurons in similar tasks. Specifically, across task conditions and trial types, we have only observed increases in CF activity (Heffley and Hull, 2019; Heffley et al., 2018; Hull, 2020). Such increases occur both for unexpected reward delivery in naive animals, and for the unexpected omission of reward when reward contingencies are reversed. These findings are consistent with previous data in both Pavlovian and operant conditioning tasks (Heffley and Hull, 2019; Heffley et al., 2018), and may suggest that CF responses reflect the type of ‘unsigned’ prediction errors that have been observed in other brain regions (Diederen and Fletcher, 2021; Roesch et al., 2012). Notably, decreases in CF activity have been reported in the context of an aversive conditioning task on omission trials (Ohmae and Medina, 2015). It therefore remains possible that the direction of CF responses to unexpected stimulus omission is context, task or region specific (Kostadinov et al., 2019).

As with previous experiments, we find that CFs respond immediately prior to licking in naive animals. In fully trained animals, however, responses to licking/reward are abolished, suggesting that they do not reflect motor output per se. These responses also do not appear to reflect motor errors, as they are also absent on CS-trials where licking occurred inappropriately. Instead, our data suggest that CF responses in this task are more closely related to expectation, similar to previous results in a reward-based operant conditioning task (Heffley et al., 2018). In line with this idea, we found that CF responses to licking re-emerged during reward contingency reversal experiments when expectation was altered. Moreover, we observed a reduced but incomplete reduction of lick-associated responses under conditions of abbreviated training when expectation should be weaker. Alternatively, it has been suggested that learned CF responses in rewarding tasks can instead represent a signal for ‘readiness to act’, occurring preferentially to learned stimuli that indicate a required behavior (Bina et al., 2022). However, because our data indicate that CFs can produce equivalent responses to similar stimuli that do and do not drive behavioral responses, the learned CF responses in our task do not appear to be directly coupled to behavior.

Motivational salience is another closely related concept that been described as part of the phasic sensory response of dopamine neurons. In short, this concept describes the property of an alerting signal to prime an animal for either action or attention toward additionally informative stimuli (Bromberg-Martin et al., 2010; Schultz, 2016). Similar to the CF responses we have measured, DA neurons can reflect the motivational salience of stimuli that are unpredictive of reward when they are presented within a rewarding context. However, in contrast with our results, such responses should also be lost as the lack of direct reward association becomes apparent. Therefore, future work will be required to test whether and how CFs in lobule simplex distinguish motivational value and salience.

Finally, our data reveal that CFs can flexibly adapt their responses to stimuli based on changing reward contingencies. Remarkably, these adaptations occurred rapidly, with responses to previously rewarded stimuli falling dramatically across the first session of rule reversal. Moreover, unexpected reward delivery in these sessions produced a significant response. However, CF responses to the new reward-associated stimulus in these experiments did not develop during the first session. Thus, it appears that extinction learning occurs more rapidly than acquisition for CF responses. While we are not able to measure CF responses across each day of training for these extended experiments using calcium imaging due to potential tissue damage and degradation of fluorescence from continued re-imaging, future experiments using alternative approaches will be important to test exactly how CF responses are transferred between unexpected reward and reward-predictive sensory stimuli.

Together, these data provide evidence that CF responses during reward-related learning share many similarities to those of canonical midbrain dopaminergic neurons. Specifically, they are context dependent, can develop learned responses that reflect reward-associated sensory input, and can be flexibly updated according to changing reward conditions. However, notable differences such as the lack of clear signed prediction errors also indicates that CF responses do not directly mimic those of midbrain DA neurons. Based on recent evidence of direct connections from the cerebellum to DA associated striatal circuits (Carta et al., 2019; Washburn et al., 2024), future experiments will be necessary to understand how the unique features of cerebellar CF responses contribute to learning and behavior in reward-based tasks that involve both brain regions.

## Methods

### Experimental animals

All experimentation using animals was performed under the approval of Duke University Animal Care and Use Committee. All experiments were performed during the light cycle in adult mice (> P60) of both sexes, randomly selected from breeding litters. All mice were housed in a vivarium with 12-hour light/dark cycles in cages with 1-4 mice. Imaging experiments were performed using Tg(PCP2-Cre)3555Jdhu mice (Jackson Labs, 010536; n = 14). All mice included in this study participated in imaging and behavioral experiments.

### Surgical procedures

3-12 hours before surgery, animals received dexamethasone (subcutaneous; 3 mg/kg). Surgical procedures were performed under anesthesia, using an initial dose of ketamine/xylazine (intraparietal; 50 mg/kg and 5 mg/kg) 5-10 minutes before surgery and sustained during surgery with 1-2% isoflurane (Paterson No. 07-893-1389). Toe pinch and breathing rate were used to monitor anesthesia levels throughout surgery. Body temperature was maintained using a heating pad (TC-1000 CWE). Custom-made titanium head plates (HE Parmer) were secured to the skull using Metabond dental cement (Parkell). A 3-mm diameter craniotomy was made over lobule simplex of the right cerebellum approximately 1.4-mm lateral and 2.8-mm posterior to lambda.

Mice were injected (WPI UMP3) with pGP.AAV.CAG.FLEX.jGCaMP7f.WPRE (Addgene 104496, initial titer = 1.5 x 10^13). Approximately 270 nL virus diluted 1:12 in aCSF was injected at 2 sites in rostral dorsal lobule simplex (at depths of 200 um [45 nL/min], 300 um [60 nL/min], and 600 um [60 nL/min]) and approximately 225 nL virus diluted 1:12 in aCSF was injected at 2 additional sites in caudal dorsal lobule simplex (at depths of 200 um [45 nL/min] and 300 um [60 nL/min]). The additional depth at the first two sites was designed to target white matter tracts to introduce the virus to a large portion of PC axons to support expression from superficial injections. After each injection at each depth, the pipette tip was left undisturbed for 3 minutes to allow for viral diffusion in an attempt to limit disruption around the site when moving to the next depth. After the final injection at a site, the pipette was left undisturbed for 5 minutes before being slowly removed and repositioned for the next injection.

The dura was kept moist with aCSF and surgical foam (Ethicon 1972) to protect the brain surface during the injection procedure. A glass cover slip, consisting of two 3-mm and one 5-mm diameter glass discs (Warner Instruments No. 64-0720 and No. 64-0700, respectively) bonded together using index matched adhesive (Norland No. 7106), was secured in the craniotomy using Metabond dental cement. A custom-made support was used to keep the glass window against the brain surface as the Metabond cured.

Buprenex (subcutaneous; 0.05 mg/kg) and cefazolin (subcutaneous; 50 mg/kg) were administered following surgery. Three additional doses were administered in 12-hour increments following the initial dose to aid recovery. Imaging was performed no sooner than 14 days after injection to allow time for adequate viral expression and to provide sufficient time to monitor craniotomy health.

### Behavior

Animals were water deprived for 3 days, after a minimum of 7 days post-surgery. Animals were then habituated to head restraint for a minimum of 4 days before experimentation began. During behavioral training, animals were head-fixed and placed in front of a computer monitor and reward delivery tube. A piezoelectric sensor (C.B. Gitty, 41 mm ‘jumbo’ piezo, No. 50-005-02) was used to measure animal movement, and a custom contact lick sensor was used to detect licks. Animals were trained to discriminate between two cues, one which predicted the delivery of a saccharine reward (CS+, 0.01M) and another which predicted no reward (CS-). A mean luminance grey screen persisted throughout the session, including inter-trial intervals (ITI, 12-18 s). During presentation of visual cues, a moving grating replaced the grey screen. The CS+ (Vis+) in both experiments was a high-contrast horizontal grating drifting rightwards for 767 ms. Termination of CS+ coincided with reward delivery in all trials. In all interleaved experiments, CS+ and CS-presentations were randomly interleaved so that cue presentation on previous trials carried no information about cue presentation in future trials. The number of CS+ and CS-trials were therefore approximately equal across all imaging and training sessions. For blocked Vis+/Vis-experiments, CS+ and CS-presentation was separated into two distinct blocks of 60 trials. Block A consisted of 60 CS+ trials followed by 60 CS-trials and Block B consisted of 60 CS-trials followed by 60 CS+ trials. Mice pairs were randomly assigned Block A or Block B for the first blocked session and were assigned the opposite block design for the second session. This was done to control for any effects of block order on neuronal response to stimuli.

In Vis+/Vis-experiments, the CS-was a high-contrast vertical grating drifting upwards for 767 ms. All characteristics of the CS-were matched with those of the CS+ (e.g. luminance, speed of grating, contrast) apart from the orientation and direction of movement. In Vis+/Aud-experiments, the visual grating was replaced by an auditory tone (1kHz, avg. 67 dB) presented for 200 ms. For mice that underwent both Vis+/Aud- and subsequent reversal training (Aud+/Vis-), the duration of the auditory tone (1 kHz, avg. 67 dB, 767 ms) was adjusted such that the visual and auditory cues were presented for the same duration. We measured no differences in behavioral or neural responses associated with auditory cue duration, so data were collapsed for final analyses.

For Vis+/Vis-experiments, training sessions lasted for 49.5 ± 2.0 min and consisted of 124.2 ± 7.2 trials (n = 171 sessions). Imaging sessions lasted for 48.3 ± 1.9 min and consisted of 120.7 ± 2.5 trials (n = 14 sessions). For Vis+/Aud-experiments, training sessions lasted for 53.0 ± 2.1 min and consisted of 127.2 ± 4.1 trials (n = 96 sessions). Imaging sessions lasted for 49.0 ± 0.2 min and consisted of 124.0 ± 0.5 trials (n = 11 sessions; 3 mice with 1 pre-learning and 1 post-learning session, and 5 mice with 1 post-learning session). For Aud+/Vis-, training sessions lasted for 54.9 ± 1.8 min and consisted of 138.6 ± 4.6 trials (n = 51 sessions). Imaging sessions lasted for 48.9 ± 0.4 min and consisted of 124.3 ± 0.9 trials (n = 10 sessions; 5 mice with 2 sessions).

Behavioral parameters, including cue presentation, reward delivery, and lick responses, were monitored using MWorks (http://mworks-project.org) and custom software written in MATLAB (Mathworks)(Heffley and Hull, 2019). Learning was defined by stabilization of reaction times and hit rates, with required reaction times for CS+ trials less than 900 ms and a hit rate above 95% and required miss rates for CS-trials greater than 90%. Across animals, in Vis+/Vis-experiments, 28.5 ± 4.8 training sessions, in Vis+/Aud-experiments, 12.0 ± 1.2 training sessions, and in Aud+/Vis-experiments, 10.3 ± 0.6 training sessions were required to meet these criteria.

### Calcium Imaging

Two-photon imaging was performed with a resonant scanning microscope (Neurolabware) using a 16x water immersion objective (Nikon CFI75 LWD 16xW 0.80NA). The space between the imaging window and objective was filled with a polymer-stabilized immersion solution (MakingCosmetics, 0.4% Carbomer 940). A Ti:Sapphire laser tuned to 920 nm (Spectra Physics, Mai Tai eHP DeepSee) was raster scanned via a resonant galvanometer (8kHz, Cambridge Technology) at a frame rate of 30Hz with a field of view of 1030 x 581 um (796 x 264 pixels). Data were collected through a green filter (510 ± 42 nm band filter (Semrock)) onto GaSaP photomultipliers (H10770B-40, Hamamatsu) using Scanbox software (Neurolabware). Experiments were aborted if significant z-motion was evident during recording.

A total of 14 mice were selected for imaging experiments across both experiments. In these experiments, the range of contributing dendrites and mice to each experiment was as follows: Vis+/Vis-experiments: pre-learning, 22-142 dendrites, 6 mice; post-learning, 24-84 dendrites, 6 mice, blocked sessions 13-90 dendrites, 6 mice; Vis+/Aud-experiment: pre-learning, 61-142 dendrites, 3 mice; post-learning, 37-139 dendrites, 8 mice; Aud+/Vis-experiments: pre-learning, 51-138 dendrites, 5 mice; post-learning, 37-134 dendrites, 5 mice.

## Data analysis and statistics

### Behavior

Analysis was performed with custom MATLAB code. Reaction times were defined by the first lick of a lick burst between 200-1500 ms after cue onset. Lick bursts were defined as a sequence of at least 3 licks within a 300 ms window. Hit rates were defined as the percentage of trials with lick bursts within 1500 ms from cue onset, or 733 ms following reward delivery on Vis+ or Aud+ trials. Correct reject rates are defined as the absence of a lick burst during the same window on Vis- or Aud-trials.

### Two-photon imaging

Motion in the X and Y planes were corrected using sub-pixel image registration. To isolate signals from individual Purkinje cell (PC) dendrites, we used a post-hoc processing pipeline consisting of principal component analysis (PCA) followed by individual component analysis (ICA)(Mukamel et al., 2009). Final dendrite segmentation was achieved by applying a gaussian filter to and thresholding pixels of individual components. A binary mask was created by combining highly correlated pixels (correlation coefficient > 0.90) and removing overlapping regions between segmented dendrites. Z-planes were chosen to limit PC soma appearing in the field-of-view, however, in the case that PC soma was present, image segmentation avoided extracting soma as regions-of-interest. Fluorescence changes (ΔF) were normalized to a window of baseline fluorescence (F) 4000 ms preceding trial initiation for each trial.

To extract events corresponding to Cspks, the raw fluorescence trace was passed through a deconvolution pipeline (OASIS: Online Active Set Method to Infer Spikes (Friedrich et al., 2017)). Cells with activity that failed to return to baseline within 33 seconds (average duration of 4 trials) or cells whose spike rate was 3 standard deviations below the population average, were classified as “corrupt” and removed from analysis. For all analysis, the post-cue period is defined as 767 ms between cue and reward delivery, and post-reward period is defined as 767 ms after reward delivery. Statistical tests for significant Cspk rates compared the average of a 3-frame window centered around the peak amplitude within post-cue and post-reward periods to a 767 ms window before cue onset (baseline). Significance for motor-aligned responses was calculated by comparing the amplitude of a 3-frame average centered around the peak response prior to motor onset with the average peak response of the baseline period.

Cspk rates represent the session-averaged number of spikes for each cell. Data are first aligned to onset of cue presentation or motor event and then mean and standard error of the mean (SEM) are calculated to represent cue or motor aligned activity. To determine response preference for individual cells (**Supp. Fig. 1A**), we determined the modulation index of peak session-averaged activity of a neuron in response to cue 1 and cue 2 presentation (modulation index = (Cspk rate_Vis-_-Cspk rate_Vis+_)/( Cspk rate_Vis-_ + Cspk rate_Vis+_). Cells with a modulation index greater than 0.33 were considered cue 1-preferring cells, those with a modulation index lower than -0.33 were considered cue 2-preferring cells, and cells with modulation indices between -0.33 and 0.33 were considered non-preferring. Finally, cells with peak session-averaged activity below 1.0 Hz were considered non-responsive for this comparison.

### Statistics

Data are presented as mean ± SEM, unless noted otherwise. Statistical significance was defined as P < 0.05. No statistical tests were performed to predetermine sample sizes. Data collection and analysis were not performed blind to the experimenters, but both data collection and analysis relied on automated measurements and were uniformly applied to all animals across each experiment.

## Acknowledgements

This work was supported by grants from the following NIH institutes: NINDS 5R01NS096289 (CH), NINDS R01NS112917 (CH), NINDS R01-NS128054 (CH). The funders had no role in study design, data collection and analysis, decision to publish or preparation of the manuscript. We thank Wenjuan Kong for technical support on this project, and Dr. Lindsey Glickfeld for comments on the manuscript.

## Author Contribution Statement

C.H., C.V. and M.M. designed experiments. C.V. and M.M. conducted experiments and analyzed data. C.H. and C.V. wrote and edited the manuscript.

## Data Availability

Due to the large size of these datasets, data that support the findings of this study are available from the corresponding author upon request.

## Code Availability

Code that support the findings of this study are available at: https://github.com/Glickfeld-And-Hull-Laboratories/ImagingCode-Glickfeld-Hull/tree/master/carlo/Vignali_flexibleCFResponses

---

**Supp. Figure 1.**
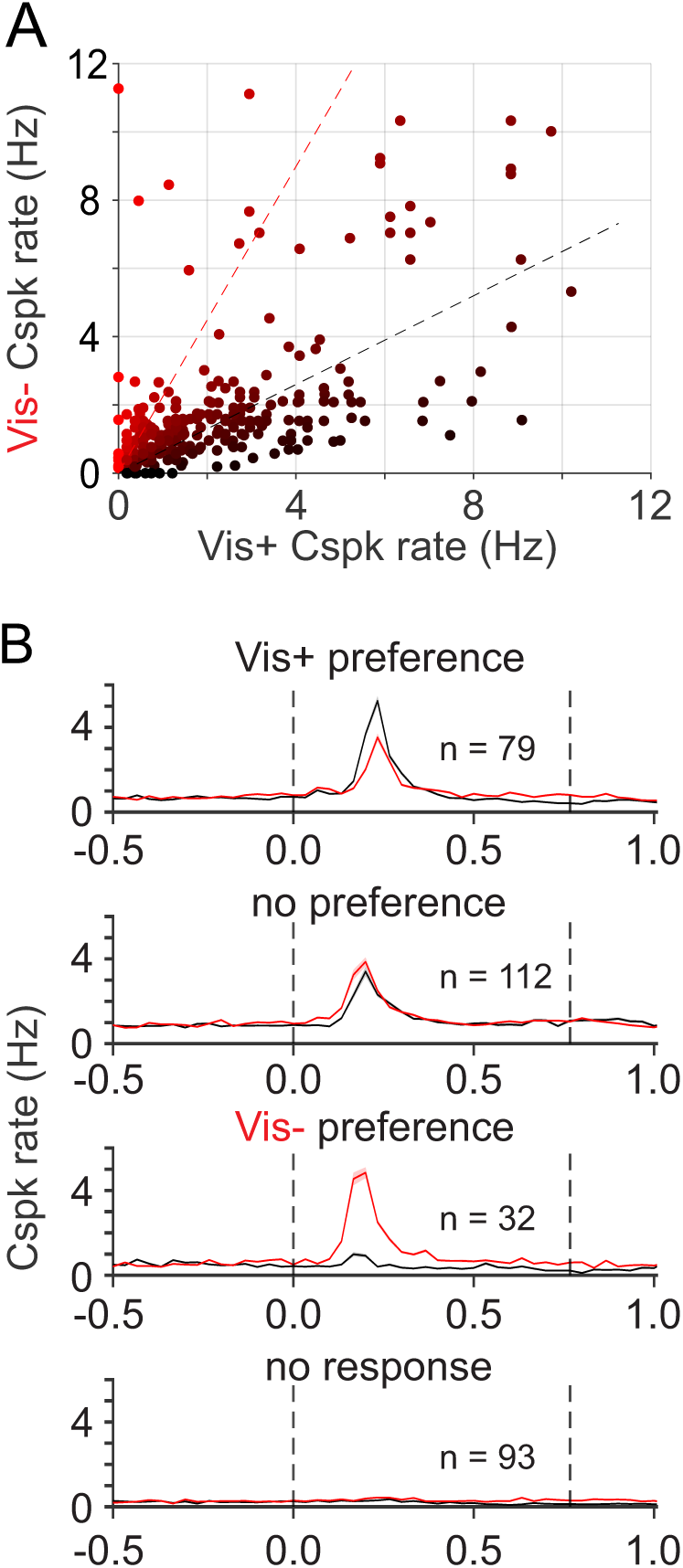
Individual CF inputs are not specific to Vis+ or Vis-. **A)** Summary of individual neuron Cspk responses to Vis+ and Vis-presentation. Red and black dashed line represents cutoff for neurons characterized as Vis- and Vis+ preferring, respectively. Cells between the two lines were classified as having no preference. Cells with mean complex spike rate below 1Hz were classified as nonresponsive. **B)** PSTHs of Cspk rates for cells based on classification in A). n = cells. All data are presented as mean +/- SEM.

**Supp. Figure 2.**
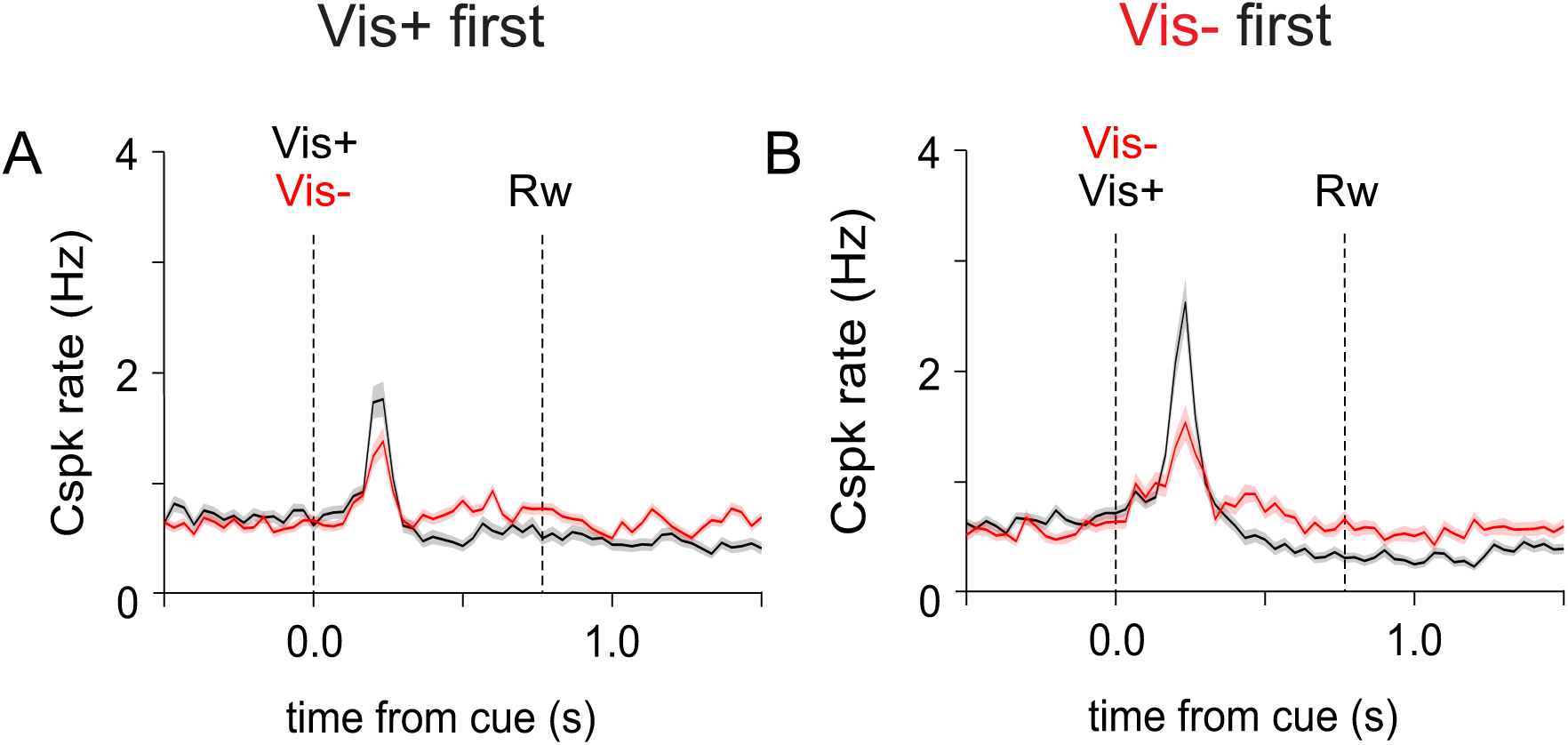
CFs preferentially respond to Vis+ on blocked trials regardless of order. **A)** PSTH of Cspk rates for trial block sessions where the Vis+ block (black) was presented before the Vis-block (red). **B)** Same as A) for sessions where Vis-block was presented first. All data are presented as mean +/- SEM.

